# D-CoMEx: A Unified Approach to Identify Phenotype-specific miRNA Biomarkers Related to Diseases

**DOI:** 10.1101/2025.06.10.658948

**Authors:** Tulika Kakati, Dhruba K Bhattacharyya, Jugal K Kalita

## Abstract

Profiling of miRNAs is important to understand how they regulate biological pathways and contribute to diseases. With current technological advancements leading to capture of voluminous biological data, it is interesting and challenging to explore new methods that can discover disease-data associations and provide better insights into the underlying mechanisms of diseases. Such understanding may also help prevent severe pathogenesis of the disease and contribute to knowledge required for effective drug development. This paper proposes a unified approach, D-CoMEx, to mine miRNA biomarkers from dysregulated pathways associated with Parkinson’s disease (PD) and Breast Cancer (BC). The approach integrates a co-expression network (CEN) module extraction technique with a differential analysis method to capture interactions between miRNAs and dysregulated pathways. Our method identifies more statistically and biologically significant dysregulated pathways compared to DCGL, MODA, and DiffCoEx. We present three evidence-backed recommendations supporting strong associations between miRNAs and dysregulated pathways, demonstrating that D-CoMEx extracts biologically relevant miRNA biomarkers linked to disease progression.

## 1 Introduction

An effective and efficient computational method can significantly help in better understanding of disease-data associations. The analysis of such associations can help uncover functionality hidden within copious biological data (such as gene expression, protein-protein interaction, and miRNA data) and elucidate mechanisms related to human diseases. Recent generation and accumulation of data through microarray and next generation sequencing (NGS) technologies have accelerated research in the exploration of disease-data interactions. Many computational methods and tools have been proposed to help identify disease related biomarkers in sources of biological data specific to a disease. For example, Wang et al. proposed a method which can identify miRNA biomarkers and discover their functionality relevant to a disease[1]. Kakati et al. proposed a co-expression module extraction technique, which can mine biomarkers related to AD from genes which have high semantic similarity but low expression similarity with the core gene of a module[2]. Another class of computational methods[3] integrates techniques related to gene expression and proteinprotein interactions. Such an integrated approach can lead to better interpretation of disease-data and identification of biomarkers related to a disease.

In the recent past, several significant computational methods and tools[4, 5, 6, 7, 8] have been developed to identify and analyze co-expressed modules in terms of functional relationships with a specific disease. A class of methods[9] and[10] have been proposed by Le et al. to identify the miRNA biomarkers from heterogeneous networks of miRNAs and their gene targets using miRNA-target gene interaction databases such as miRWalk, and TargetScan. The concept of dysregulated pathways[3] characterizes the impairment in regulation of gene-gene interactions taking into account disease and control samples. Dysregulated pathways overcome the limited descriptive power of individually dysregulated genes by considering genes as members of biologically or functionally similar network modules, not individual entities. The main limitations of any CEN method include, it cannot find causal entities and cannot detect the changes in co-expression patterns across different conditions or tissues because for the same pair of genes under different conditions or tissues, the correlation measure may show different correlation. In other words, regular coexpression or differential analysis cannot identify the underlying differences between control and disease conditions[11]. An approach, such as differential co-expression analysis can identify changes in expression patterns between conditions and can help biologists identify the potential regulators of a particular disease. Most methods, such as DICER[12], DiffCoEx[13], DCGL[14], MODA[15], and METADCN[16] identify differentially co-expressed genes and detect modular changes in the genes across conditions. Another method DiffCoMO[17] uses multi-objective approach to mine differentially co-expressed gene modules across two different stages of HIV-1 progression. However, to the best of our knowlegde, none of the methods use miRNA-mRNA jointly profiled datasets to identify the miRNA-disease interactions. In THD-Module Extractor[2], Kakati et al. use a unique feature of core gene to identify the co-expressed modules based on the expression discriminative behavior of core gene and with the border genes(s). The core gene of a CEN module with the highest connectivity to other genes is found to have high biological significance with border genes of the module. Therefore, we find the analysis of the differentially co-expressed genes with respect to the core gene as a key aspect in identifying the potential biomarkers of a disease. Table 1 shows the key features of the six differential co-expression analysis techniques.

**Table 1:**
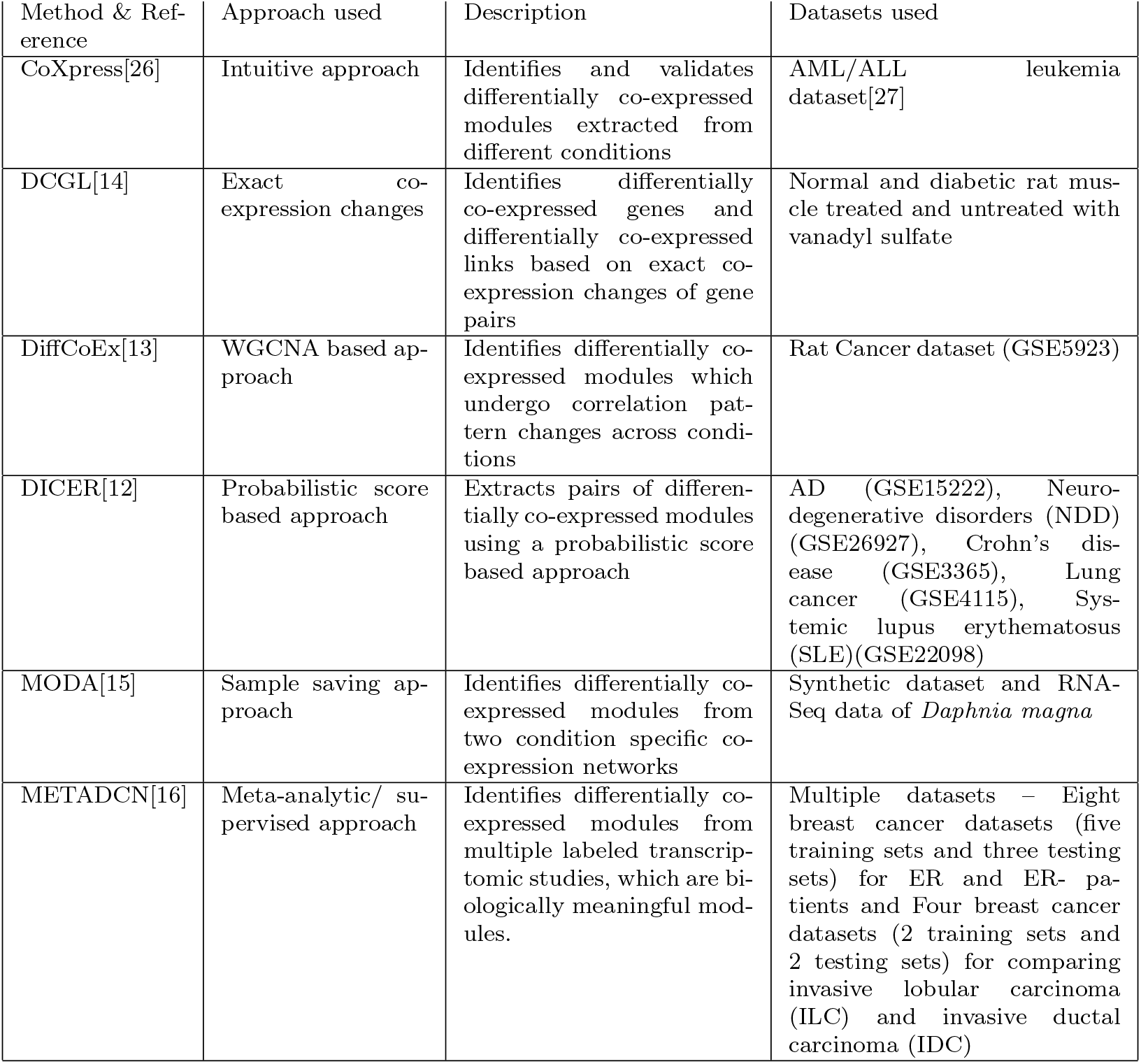
Key features of differential co-expression analysis techniques.

This paper introduces a novel approach by fusing a THD-Module Extractor with a differential analysis technique, to identify potential miRNAs from dysregulated pathways associated with Parkinson’s Disease (PD) and Breast Cancer (BC). The differentially co-expressed mRNAs and miRNAs extracted from mRNA-miRNA expression profiles are mapped onto pathways collected from the Biocarta[18], Kegg[19], and Reactome[20] databases using ToppGene Suite[21], Panther[22], and MiRSystem[23]. These pathways, known as dysregulated pathways are condition-specific and undergo perturbation during pathogenesis of the disease. This analysis suggests that correlation between dysregulated pathways can be used to infer novel pathways for a disease unknown to existing literature. The mRNA/miRNA modules extracted from mRNA-miRNA datasets are validated considering both statistical and biological significance using web-based tools such as TopoGSA[24], ToppGene Suite[21], Panther[22], and MiRSystem[23].

In contrast to the work in[3], the dysregulated pathways found using our proposed approach are further analyzed to identify miRNA biomarkers and predict the functional similarity of the miRNAs related to a disease. miRNAs modulate the expression of mRNAs of target genes, which in turn alter biological pathways and play significant role in host-pathogen interactions[25]. The miRNAs regulating the dysregulated pathways are either up-regulating or down-regulating during the progression of a disease. Modeling dysregulated pathways across disease specific stages can fundamentally alter the perception of how pathways or cellular networks behave during pathogenesis of diseases. Furthermore, identification of such gene targeting miRNAs from dysregulated pathways can help gain insights into biomarkers related to a disease and eventually help in effective drug development.

## 2 METHODS

We implement the proposed method as an *R* package called D-CoMEx (Differential Co-expressed Module Extraction) to extract differentially co-expressed modules from mRNA-miRNA expression data combining a co-expressed network module extraction technique and a differential analysis technique. The work-flow of the proposed approach is shown in Figure 1. The steps in the framework are detailed in the following subsections.

**Figure 1:**
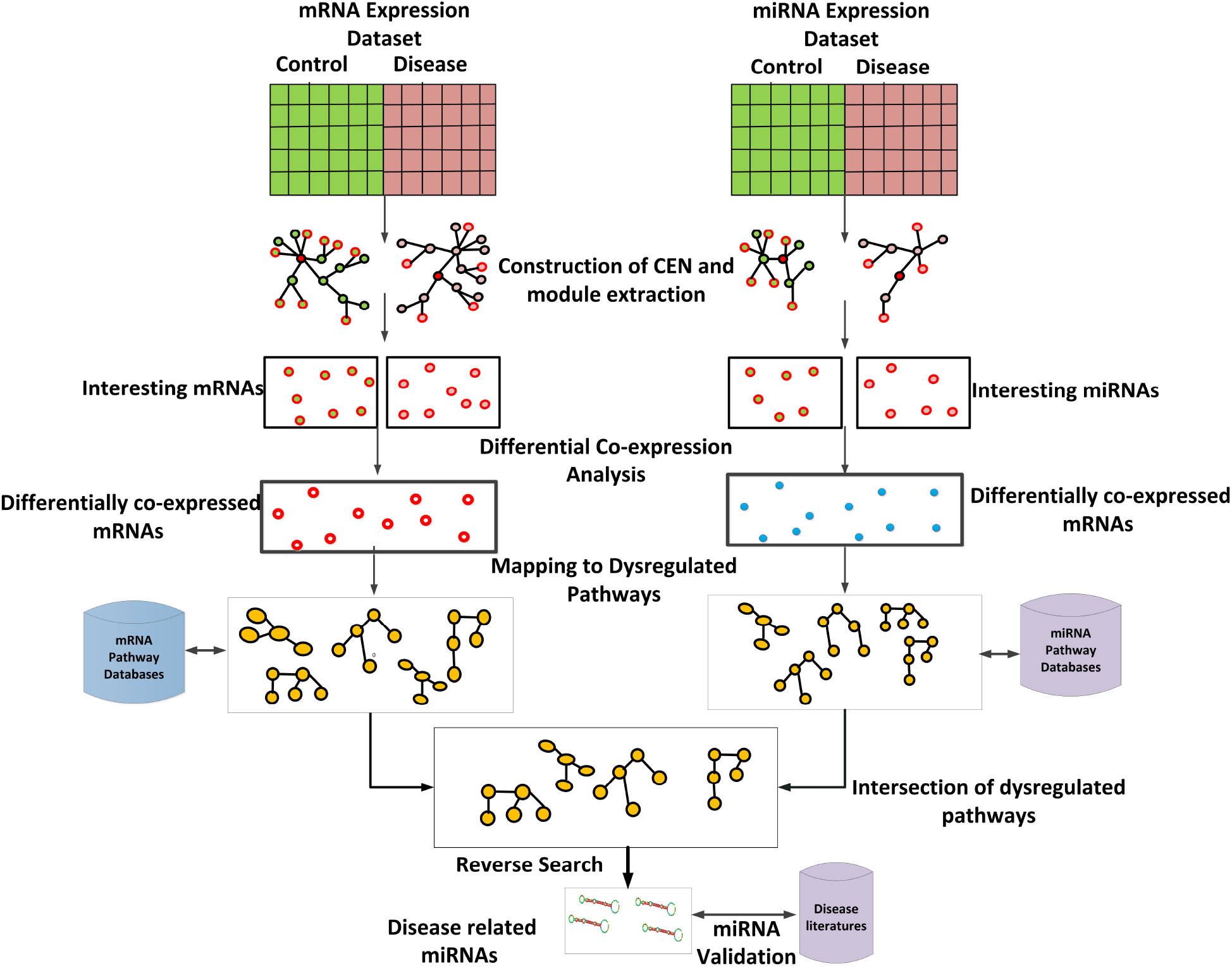
Workflow of the current proposed approach. This method combines CEN module extraction and differential analysis techniques. The co-expressed modules are extracted from control and disease states of both mRNA and miRNA datasets. Differential co-expression analysis is performed on the interesting mRNAs and miRNAs of the co-expressed modules extracted from both expression profiles. The differentially co-expressed mRNAs and miRNAs are mapped to pathways using the ToppGene Suite[21], Panther[22], and MiRSystem[23], respectively. From the common dysregulated pathways, we filter out the miRNAs using a reverse search technique in DIANA (mirPath v.3)[28]. miRNAs regulating these dysregulated pathways are found to promote the progression of the diseases.

### 2.1 Data used and preprocessing

For this study, we download two jointly profiled mRNA-miRNA datasets (GSE77668 and GSE19783) from the Gene Expression Omnibus (GEO) database[29] with both mRNA and miRNA expressions in control (or normal) and disease (or tumor) patients. The details of the datasets (Table 1 of Supplementary file) and the preprocessing activity of each dataset are given in the Supplementary file.

### 2.2 Extraction of co-expressed modules using THD-Module Extractor

We use THD-Module Extractor[2] to extract co-expressed modules from both control and disease conditions of mRNA and miRNA expression profiles. The THD-Module Extractor accepts one parameter, minimum neighborhood threshold (*ρ*). CEN is a graph based representation of pairwise relationships among genes or gene products, where genes are represented by nodes and relationships between the nodes are represented by edges[30]. The method constructs a CEN from a gene expression dataset and extracts co-expressed modules of genes with high expression similarity among them using the similarity measure called SSSim[31]. In this method, a gene in a module is called the core gene if it is the starting gene of the module, which has the highest number of connected nodes and satisfies the minimum number of neighbors threshold. The core gene of a module does not belong to any other modules of the CEN.

The process of module extraction progresses gradually by decreasing *δ* by a constant factor (*β*) of 0.05 from upper bound expression similarity threshold (*δ*)=0.95 to lower bound expression similarity threshold (*δ*)=0.35. In our study, we choose the minimum neighborhood threshold *ρ* as 250 heuristically throughout the experiment to optimize the results and avoid the generation of modules with low cardinality. This is evidenced by the graph given in Figure 1 in the Supplementary file.

In this work, we use Resnik measure[32] to find semantic similarities between the mRNAs and the genes targeted by miRNAs of the co-expressed modules of each expression profile, since Resnik measure is more appropriate to measure semantic similarity of a pair of genes. From the mRNA coexpressed modules, we consider the border mRNAs with high semantic similarities as interesting. On the other hand, from miRNA co-expressed modules, we consider the border miRNAs with maximum average semantic similarities shared by their target genes as interesting.

### 2.3 Differential analysis of interesting co-expressed modules

The mRNAs and miRNAs in co-expressed modules extracted from both control and disease conditions have condition specific high connectivity and high semantic similarity among them. We apply a differential co-expression analysis technique to study the perturbation of the interesting mRNAs, i.e., genes and miRNAs across normal conditions and disease conditions[33]. The set of co-expressed mRNAs or miRNAs identified in one set of conditions, which undergo changes in expression behavior across another set of conditions are defined as differentially co-expressed mRNAs or miRNAs. The pathways mapped to these differentially co-expressed mRNAs and miRNAs are called as dysregulated pathways.

1. A group of genes or mRNAs *G*^*′*^=*{g*_1_, *g*_2_, …, *g*_*m*_*}* is referred to as differentially co-expressed if they are co-expressed in one set of conditions and undergo significant expression or behavioral change in another set of conditions.
2. An mRNA, i.e., a gene *g*_*i*_ or miRNA *m*_*i*_ is defined as interesting if (i) it lies on the periphery of a co-expressed module and (ii) it has high semantic similarity with the core gene of the module.

Suppose, co-expressed module *M*_*i*_ has a set of genes *{g*_1*c*_, *g*_2*c*_, *g*_3*c*_, *g*_4*c*_*}* across control conditions and module *M*_*j*_ has a set of genes *{g*_1*d*_, *g*_2*d*_, *g*_3*d*_, *g*_4*d*_*}* across disease conditions. Now, to find the differentially co-expressed genes from these modules, we follow the following steps.

1. For each pair of genes (*g*_*ic*_, *g*_*jd*_), if *V*_*i*_, *V*_*j*_ are two matrices of expression values across control and disease conditions, we find Cov (*g*_*ic*_, *g*_*jd*_) = *p*_*ij*_/(len(*V*_*i*_)-1), where 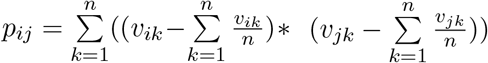 and *n* is the total number of samples in control and disease conditions.
2. If Cov (*g*_*ic*_, *g*_*jd*_) is negative then the genes or miRNAs covary in opposite direction. This means that if *g*_*ic*_ is up-regulating in control stage then *g*_*jd*_ is down-regulating in disease stage. This signifies that there exists variance in their expression profiles across control and disease conditions.
3. In order to find the strength of this variance, we find the correlation between the genes (*g*_*ic*_, *g*_*jd*_) as Corr (*g*_*ic*_, *g*_*jd*_) = Cov (*g*_*ic*_, *g*_*jd*_) / (std(*V*_*i*_)*std(*V*_*i*_)).
4. If Corr (*g*_*ic*_, *g*_*jd*_) *<* 0.75, then *g*_*ic*_ and *g*_*jd*_ have low covariance across control and disease stages and therefore they are differentially co-expressed.

The common dysregulated pathways mapped from both mRNAs and miRNAs as well as the numbers of total and common dysregulated pathways are specified in Supplementary file. The common dysregulated pathways, which echo the interactions among mRNAs and molecules for a particular biological process, are also regulated by the differentially co-expressed miRNAs. The validation of the miRNAs regulating dysregulated pathways provides insights into the miRNAs associated with these deadly neurodegerative diseases. For validation of the miRNAs regulating the dysregulated pathways, we have downloaded the experimentally validated miRNA-mRNA data for human from the miRWalk 2.0 database[34]. The differentially co-expressed miRNAs extracted using D-CoMEx, are found to target the differentially co-expressed mRNAs extracted using DCoMEx (results are given in Supplementary file).

The dysregulated pathways are both statistically and biologically interpretable and may exhibit discriminative features. Therefore, we assess the behavior of the identified dysregulated pathways mapped from mRNAs and miRNAs. We hypothesize that the miRNAs regulating the dysregulated pathways are differentially co-expressed and therefore they play vital role in pathogenesis of a disease. In turn, we assume the existence of a strong association between miRNAs and dysregulated pathways. We present recommendations in support of the association between miRNAs and dysregulated pathways, which are discussed in the Results section. We give the pseudocode for the THD-Module Extractor and D-CoMEx in Algorithms 1 and 2, respectively. The symbols used in the algorithms are listed in Table 2.

**Table 2:** Descriptions of symbols.

| Symbols | Description | Symbols | Description |
| --- | --- | --- | --- |
| $D$ | Dataset | $V$ | Co-expression Network (CEN) |
| $(g_i, g_j)$ | Gene pair or mRNA pair | $(m_i, m_j)$ | miRNA pair |
| $\delta$ | Expression similarity threshold | $\rho$ | Minimum neighborhood threshold |
| $\beta$ | Semantic similarity threshold | $g_b$ | Border gene or mRNA |
| $m_b$ | Border miRNA | $g'_b$ | Interesting $g_b$ |
| $m'_b$ | Interesting $m_b$ | $G_b$ | Set of $g_b$ |
| $M_b$ | Set of $m_b$ | $G'_b$ | Set of $g'_b$ |
| $M'_b$ | Set of $m'_b$ | $g_d$ | Differentially co-expressed $g'_b$ |
| $G'_d$ | Set of differentially co-expressed $g'_b$ | $M'_d$ | Set of differentially co-expressed $m'_b$ |
| $\delta_d$ | User defined constant for $g_d$ | $A_{ij}$ | Adjacency matrix |
| $M$ | Co-expressed module | $nNeighbor(g_i)$ | Number of neighboring genes of gene $g_i$ |
| $S_{ij}$ | Semantic similarity correlation matrix | $Exp(g_{id})$ | Expression of gene $g_i$ for a disease sample $d$ |
| $Exp(g_{ic})$ | Expression of gene $g_i$ for a control sample $c$ | $V_i, V_j$ | Row matrix with difference of expressions between disease and control samples for gene $g_i$ and $g_j$ respectively |

### 2.4 Identification of miRNA biomarkers associated with disease

A systematic study of miRNAs associated with dysregulated pathways in diseases like PD and BC would provide better insights into the progression of diseases and may help further investigation into these diseases, with an eye to generating knowledge to support effective drug development. Chen et al.[35] reviewed 20 computational methods of predicting miRNA-disease associations and summarized the issues in predicting disease related miRNAs and difficulties in constructing models to predict miRNA-disease associations. They divided the prediction models into four types, namely, machine learning based (e.g., MCMDA[36], LRSSLMDA[37], and EGBMMDA[38]), score function based (e.g., WBSMDA[39]), complex network algorithm based (e.g., PBMDA[40]), and multiple biological information based models (e.g., KBMFMDI[41]). There have also been several attempts[42, 43, 44] at discovering association between miRNAs with disease related pathways or biological functions. For example, a method described in[45], summarizes the role of miRNAs in signaling pathways in Systemic Sclerosis. Liu and Liu[46] proposed a combination method to find 13 significant differential pathways in the hippocampus of AD patients. To the best of our knowledge, this work by Liu and Liu is the only approach similar to ours, which identifies differential pathways associated to AD. They identified 13 differential pathways related to AD using mRNA and miRNA expression profiles (E-GEOD-1297, E-GEOD-5281, and E-GEOD-28146). However, Liu and Liu did not analyze differential pathways to find miRNA biomarkers regulating pathways related to a disease. In contrast, in our method, we identify the differential pathways (called as dysregulated pathways) and the corresponding miRNAs related to PD and BC, regulating the dysregulated pathways. On the other hand, methods proposed by Le et al.[9, 10] use interaction databases, such as miRWalk and TargetScan to identify the disease related miRNAs from a heterogeneous network of miRNAs and their gene targets.

#### Algorithm 1 THD-Module Extractor

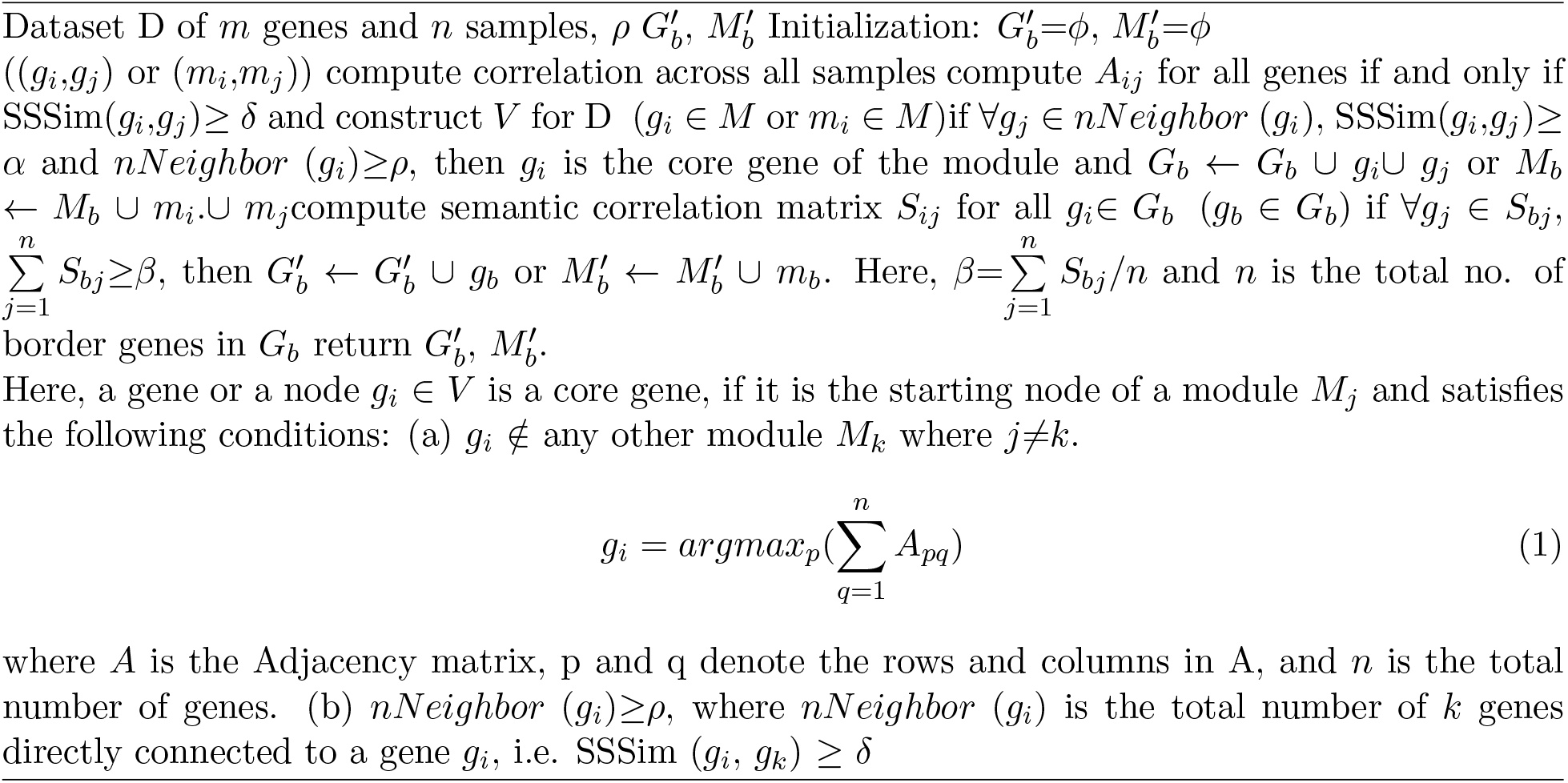

#### Algorithm 2 D-CoMEx

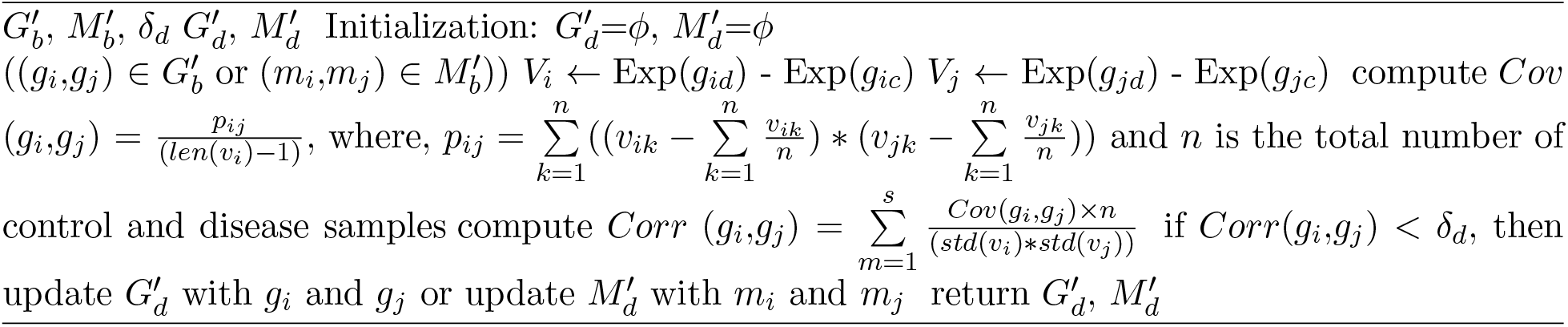

In this work, we use a reverse search approach to identify the miRNAs, which are predicted or validated experimentally to target dysregulated pathways obtained from mRNA/miRNA expression profiles. The miRNAs regulating dysregulated pathways are identified using DIANA-miRPath v3.0[28] and validated with existing literature to find their association with these deadly diseases. The potential miRNAs regulating the dysregulated pathways may help researchers predict their roles in pathology and also as potential therapeutic targets.

## 3 EXPERIMENTS AND RESULTS

We used two disease datasets, PD and BC as two case studies to check the generalizability of our method D-CoMEx in finding disease related miRNA biomarkers mapped with dysregulated pathways. In this section, we present the enrichment analysis of differentially co-expressed mRNAs and miRNAs and analysis of dysregulated pathways mapped from the differentially co-expressed interesting mRNAs and miRNAs, extracted from two disease datasets, PD and BC. We compare our results with those of DCGL[14], MODA[15] and DiffCoEx[13]. We present three recommendations in support of the association of miRNA and disease, which are explained below.

### 3.1 Enrichment analysis of differentially co-expressed mRNAs and miRNAs

In this work, we analyze and validate the differentially co-expressed mRNAs and miRNAs extracted from mRNAs and miRNAs both statistically and biologically.

In Table 3, we show the comparison of differentially co-expressed mRNAs modules extracted using D-CoMEx, DCGL,MODA, and DiffCoEx. The differentially co-expressed mRNAs extracted by our approach from mRNA expression profiles of PD and BC datasets are with high node betweenness and degree, and are statistically significant with low *p* values and *q* value in comparison to DCGL, MODA, and DiffCoEx. Node betweeness of co-expressed modules measures the centrality of a node with respect to other nodes passing through that node. Degrees of co-expressed modules measure the closeness of one node with each other in terms of the average number of edges incident on a node. The *p* value is defined as the possibility of obtaining *k* or more genes in a cluster of size *n*, associated with a specific GO term. If *f* is the number of genes in the reference list already associated with the GO term and *g* is the total number of genes in the reference list, then *p* value is formulated as

**Table 3:** Analysis of top *k* differentially co-expressed modules extracted from mRNA expression of PD and BC dataset.

| 2*Disease | 2*Methods | Module No. | Number of mRNAs | Node-betweenness | Degree | p-value | q-value |
| --- | --- | --- | --- | --- | --- | --- | --- |
| 10*PD | D-CoMEx | Module 1 | 48 mRNAs | 14,1507 | 29.45 | 2.64E-31 | 7.48E-29 |
|  |  | Module 2 | 17 mRNAs | 46,820 | 18.13 | 5.39E-10 | 4.34E-08 |
|  |  | Module 3 | 17 mRNAs | 38,235 | 16.62 | 2.48E-17 | 3.59E-15 |
|  |  | Module 4 | 18 mRNAs | 10,914 | 7.56 | 6.63E-27 | 5.90E-25 |
|  | DCGL | Module 1 | 16 mRNAs | 7,124 | 7.4 | 1.33E-28 | 6.11E-27 |
|  | MODA | Module 1 | 44 mRNAs | 5,723 | 5.74 | 3.23E-28 | 4.23E-15 |
|  |  | Module 2 | 31 mRNAs | 2,357 | 14.38 | 5.34E-14 | 2.33E-11 |
|  | DiffCoEx | Module 1 | 27 mRNAs | 5,912 | 15.38 | 2.21E-10 | 4.59E-08 |
|  |  | Module 2 | 24 mRNAs | 8,189 | 7.88 | 3.05E-27 | 3.14E-22 |
| 10*BC | D-CoMEx | Module 1 | 557 mRNAs | 19452 | 10.18 | 5.40E-10 | 6.80E-07 |
|  |  | Module 2 | 128 mRNAs | 14421 | 9.21 | 4.92E-06 | 1.54E-02 |
|  |  | Module 3 | 131 mRNAs | 16444 | 9.21 | 4.35E-05 | 1.42E-02 |
|  |  | Module 4 | 143 mRNAs | 14378 | 9.02 | 5.53E-06 | 3.39E-03 |
|  | DCGL | Module 1 | 149 mRNAs | 16527 | 10.2 | 5.07E-05 | 1.45E-02 |
|  | MODA | Module 1 | 194 mRNAs | 12499 | 8.41 | 4.22E-06 | 2.71E-03 |
|  |  | Module 2 | 260 mRNAs | 25543 | 9.6 | 7.52E-06 | 5.46E-03 |
|  |  | Module 3 | 492mRNAs | 34739 | 13.01 | 7.84E-08 | 3.89E-05 |
|  | DiffCoEx | Module 1 | 277 mRNAs | 21320 | 11.31 | 4.00E-07 | 2.77E-04 |
|  |  | Module 2 | 169 mRNAs | 9065 | 6.78 | 2.38E-07 | 1.09E-04 |
|  |  | Module 3 | 59 mRNAs | 18345 | 11.59 | 9.11E-04 | 5.75E-03 |

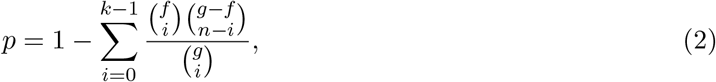

where, *p* value measures the association of GO terms with the genes. *q* value is another statistical measure, which gives the minimum false discovery rate (FDR). Lower values of *p* and *q* signify higher biological significance of the co-expressed modules. For example in Table 3, module 1 from dataset 1 with 48 mRNAs obtained from D-CoMEx outperforms among all the modules obatined using DCGL, MODA, and DiffCoEx in terms of higher values of node-betweennes and degree, and lower *p* and *q* values. Similary, among modules obatined from dataset 2, module 1 with 557 mRNAs obtained using D-CoMEx has higher topological values with node-betweenness = 19,452 and degree = 10.18 and higher statistical significances with *p* value= 5.40E-10 and *q* value= 6.80E-07 compared to all modules obtained using other three discussed methods. Therefore, the results obtained by D-CoMEx are more significant compared to those by DCGL, MODA, and DiffCoEx. In Table 4, we report the assesment of differentially co-expressed miRNA modules, extracted using the four mentioned methods in terms of *p* values of common target genes.We also validate our results using miRNA target prediction score using the MiRSystem. In Table 4, we find that among the top four modules extracted from PD and BC datasets, module 1 and module 4 with 628 miRNAs and 93 miRNAs have lower *p* value, more common target genes, higher miRNA target prediction score than all other modules extracted using other discussed methods. For example, module 1 from PD modules has *p* value= 6.81E-50, number of common target genes = 20,177, and miRNA target prediction score= 51.581. Similarly, module 3 from BC dataset has *p* value= 5.64E-07, number of common target genes = 14063, miRNA target prediction score=59.1667. A higher miRNA target prediction score signifies that the differentially co-expressed miRNAs extracted by our method exhibit more matching sequences than the discussed counterparts. Thus, we can conclude that, the differentially co-expressed miRNA modules extracted by D-CoMEx are biologically significant.

**Table 4:** Analysis of top *k* differentially co-expressed modules extracted from miRNA expression of PD and BC dataset.

| 1*Disease | 1*Methods | Module No. | Number of miRNAs | p value | Number of common target genes | miRNA target prediction score |
| --- | --- | --- | --- | --- | --- | --- |
| PD | D-CoMEx | Module 1 | 628 miRNAs | 6.81E-50 | 20,177 | 51.581 |
|  |  | Module 2 | 250 miRNAs | 1.66E-49 | 19,737 | 2.1201 |
|  |  | Module 3 | 233 miRNAs | 1.46E-43 | 19,539 | 7.1091 |
|  |  | Module 4 | 249 miRNAs | 5.69E-43 | 19,631 | 12.897 |
|  | DCGL | Module 1 | 94 miRNAs | 3.84E-20 | 17,928 | 11.0211 |
|  | MODA | Module 1 | 274 miRNAs | 7.06E-34 | 19,553 | 31.1233 |
|  |  | Module 2 | 270 miRNAs | 1.84E-45 | 19,633 | 31.605 |
|  | DiffCoEx | Module 1 | 496 miRNAs | 3.28E-38 | 20,076 | 47.1713 |
|  |  | Module 2 | 271 miRNAs | 5.43E-40 | 19,543 | 34.1212 |
| BC | D-CoMEx | Module 1 | 176 miRNAs | 3.67E-08 | 19203 | 4.38272 |
|  |  | Module 2 | 45 miRNAs | 1.64E-03 | 16477 | 22.9032 |
|  |  | Module 3 | 26 miRNAs | 5.64E-07 | 14063 | 59.1667 |
|  |  | Module 4 | 32 miRNAs | 1.48E-07 | 15662 | 39.4445 |
|  | DCGL | Module 1 | 20 miRNAs | 1.86E-07 | 1673 | 27.0476 |
|  | MODA | Module 1 | 150 miRNAs | 3.18E-02 | 16514 | 4.7651 |
|  | DiffCoEx | Module 1 | 32 miRNAs | 1.35E-02 | 16906 | 11.4516 |
|  |  | Module 2 | 55 miRNAs | 3.28E-02 | 12046 | 11.4516 |

### 3.2 Analysis of dysregulated pathways mapped from differentially co-expressed mRNAs and miRNAs

We observe discriminative characteristics of groups of differentially co-expressed mRNAs and miRNAs across the stages (control to disease) in terms of dysregulated pathways for PD and BC. It is also our observation that differentially co-expressed mRNAs included by dysregulated pathways exhibit dis-correlation among themselves across stages. Such disturbing behavior of the dysregulated pathways across the stages reveals interesting findings regarding association of a pathway for a given disease. For example, the dysregulated pathway, namely Apoptosis signaling mapped to the differentially co-expressed mRNA modules extracted from dataset 1 includes differentially co-expressed mRNAs, such as, Interleukin-1 (IL-1), Interleukin-6 (IL-6). Both IL-1 and IL-6 are included in cytokine signaling pathway and are responsible for inflammation, which accelerate the formation of lewly bodies and amyloid beta peptides, hallmarks of PD[47, 48].

We also find that most dysregulated pathways extracted from mRNA and miRNA expression profiles of the discussed datasets are common, which establishes that there must exist strong association with dysregulated pathways and differentially co-expressed miRNAs. We analyze a step further to find the miRNA biomarkers regulating these pathways and find that most miRNAs regulating the dysregulated pathways are associated with PD and BC. We present the following recommendations in support of the association between miRNAs and dysregulated pathways. *Recommendation 1:* If *P*_1_, *P*_2_, and *P*_3_ are dysregulated pathways considering control and disease conditions (states), and *P*_1_ and *P*_2_ are known to be related to a disease, it is likely that *P*_3_ is also related to the disease as well.

*Explanation:* The dysregulated pathways consist of differentially co-expressed mRNAs or miRNAs extracted from both mRNA and miRNA expression profiles. Therefore, the pathways are correlated at a particular condition (stage). In Table 5, we find the common dysregulated pathways extracted from mRNA and miRNA expression data. During pathogenesis of PD, disruptions in biological functions are controlled by dysregulated pathways, such as Wnt signaling pathway, Apoptosis signaling pathway, Parkinson’s disease, Dopamine receptor mediated signaling pathway. Modifications of these pathways induces devastating progression of PD[49, 50, 51]. We recommend that the dysregulated pathway, such as ECM signaling pathway, which is found to be correlated with the above mentioned PD related pathways may be explored further in view of their association with PD.

**Table 5:** Common dysregulated pathways extracted from mRNA and miRNA expression data and corresponding regulating miRNAs.

| Disease | Dysregulated Pathways | Differentially co-expressed miRNAs regulating dysregulated pathways |
| --- | --- | --- |
| 22*PD | Apoptosis signaling pathway | hsa-miR-195, hsa-miR-153, hsa-miR-10a-5p, hsa-miR-205, hsa-miR-128-3p, hsa-miR-433, hsa-miR-485-5p, hsa-miR-212-3p, hsa-miR-19b-3p, hsa-miR-24-3p |
|  | Parkinson's disease | hsa-miR-195, hsa-miR-153, hsa-miR-107, hsa-miR-10a-5p, hsa-miR-29b-3p, hsa-miR-590-3p, hsa-miR-205, hsa-miR-128-3p, hsa-miR-433, hsa-miR-485-5p, hsa-miR-212-3p, hsa-miR-19b-3p, hsa-miR-24-3p |
|  | Inflammation mediated by chemokine and cytokine signaling pathway | hsa-miR-195, hsa-miR-153, hsa-miR-205, hsa-miR-10a-5p, hsa-miR-212-3p, hsa-miR-19b-3p, hsa-miR-24-3p |
|  | IL-17 signaling pathway | hsa-miR-142-5p, hsa-miR-21, hsa-miR-146, hsa-miR-194, hsa-miR-182 |
|  | ECM-receptor interaction | hsa-miR-29a-3p, hsa-miR-27a-3p, hsa-miR-195-5p, hsa-miR-205-5p, hsa-miR-212-3p |
|  | Wnt signaling pathway | hsa-miR-19b-3p, hsa-miR-205, hsa-miR-29b-3p, hsa-miR-17-3p, hsa-miR-212-3p, hsa-miR-204-5p, hsa-miR-191-5p |
|  | Dopamine receptor mediated signaling pathway | hsa-miR-191-5p, hsa-miR-195, hsa-miR-205, hsa-miR-153, hsa-miR-590-3p, hsa-miR-212-3p, hsa-miR-24-3p, hsa-miR-19b-3p |
| 11*BC | Cancer pathway | hsa-miR-181, hsa-miR-155, hsa-miR-520, hsa-miR-126, and hsa-miR-200 |
|  | Cell Cycle pathway | hsa-miR-302, hsa-miR-106b, hsa-miR-106a |
|  | WNT signaling pathway | hsa-miR-34a-5p, hsa-miR-16-5p, hsa-miR-522-5p, and hsa-miR-17-5p |
|  | Hemostatis pathway | hsa-miR-335-5p, hsa-miR-124-3p, hsa-miR-34a-5p |
|  | T-Cell receptor pathway | hsa-miR-34a-5p, hsa-miR-522-5p, hsa-miR-17-5p, hsa-miR-182-5p, hsa-miR-27a-3p |

Similarly, for BC, WNT signaling pathway dramatically imbalances during pathogenesis of BC[52]. Studies demonstrate that there is seen epigenetic disturbances in the components of WNT pathways during progression of breast cancer stages. In accordance with recommendation 1, other correlated pathways such as T cell receptor may also likely to be related to BC, which has been later reported in[53].

*Recommendation 2:* If *m*_1_, *m*_2_, and *m*_3_ are differentially co-expressed miRNAs considering control and disease conditions, and *m*_1_ and *m*_2_ are known to be related to a disease, it is likely that *m*_3_ is related to the disease as well.

*Explanation:* In Table 5, we find that for PD, miRNAs, regulating Parkinson’s Disease pathway, such as hsa-miR-195, hsa-miR-153, hsa-miR-107, hsa-miR-10a-5p are differentially co-expressed and hsa-miR-195, hsa-miR-107 are related to PD[54]. Therefore, in accordance with Recommendation 2, hsahsa-miR-153[55] and hsa-miR-10a-5p[56] are also potential biomarkers of PD.

*Recommendation 3:* If *m*_1_ and *m*_2_ are differentially co-expressed miRNAs and *m*_1_ regulates pathway *P*_1_, it is likely that *m*_2_ also regulates *P*_1_ and further analysis of *m*_2_ is recommended. The same holds for the pathways regulated by *m*_2_.

*Explanation:* From Table 6, miRNA, hsa-miR-34a negatively regulates WNT signaling pathway[57] and hsa-miR-34a is found to be differentially co-expressed with hsa-miR-16-5p. Therefore, biologists can explore if miRNA, hsa-miR-16-5p may also regulate WNT signaling pathway in accordance with Recommendation 3.

**Table 6:** miRNAs regulating dysregulated pathways and related to PD and BC.

| Disease | miRNAs | miRNA characteristics |  |
| --- | --- | --- | --- |
|  |  | Up-regulation | Down-regulation |
| 5*PD | hsa-miR-195, hsa-miR-153, hsa-miR-10a-5p, hsa-miR-205 | ✓ |  |
|  | hsa-miR-128, hsa-miR-433, hsa-miR-485-5p, hsa-miR-132-5p, miR-212-3p, hsa-miR-19b-3p, hsa-miR-24 |  | ✓ |
| 5*BC | hsa-miR-130a, hsa-miR-205, hsa-miR-29b, hsa-miR-155 | ✓ |  |
|  | has-miR-34a, hsa-miR-10b, hsa-miR-145, hsa-miR-221, hsa-miR-222 |  | ✓ |

### 3.3 Comparison of D-CoMEx with existing differential co-expressed techniques

In the recent past, many computational techniques have been used to identify differentially coexpressed mRNAs or genes from microarray and RNA-seq datasets. We compare the biological significance of dysregulated pathways extracted from mRNA expression profile of the PD and BC datasets with the pathways mapped from the differentially co-expressed mRNAs extracted using three existing approaches, DCGL[14], MODA[15] and DiffCoEx[13]. The dysregulated pathways extracted using our proposed approach D-CoMEx have lower *p* values, lower FDR (B&H) values and more number of annotated genes. In MODA[15], differentially co-expressed networks are constructed from both control and disease conditions and condition specific modules are extracted from each of the networks. On the other hand, DiffCoEx[13] extracts modules undergoing correlation pattern changes across different conditions. The dysregulated pathways mapped from the best differentially co-expressed mRNAs are found to be related to PD. Therefore, these dysregulated pathways are validated in terms of *p* values, the number of annotated genes, and FDR (B&H) values using the ToppGene Suite[21]. Lower the values of *p* and FDR, the more significant is the result.

Tables 5 and 9 of Supplementary file describes the dysregulated pathways, which are known to be associated with PD and BC, respectively. Table 7 shows that the dysregulated pathways mapped from differentially co-expressed modules of PD mRNAs are more biologically significant compared to those of DCGL, MODA, and DiffCoEX results in terms of *p*, FDR (B&H) values, and number of annotated genes. For instance, the considered mRNA module from D-CoMEx and MODA has 44 number of mRNAs, but the dysregulated pathways mapped from the mRNA module of D-CoMEx are more significant with *p* value= 7.980E-18, FDR (B&H) value=6.368E-15, and number of annotated mRNAs=270. Thus, D-CoMEx outperforms the state-of-the-art methods in terms of both statistical and biological significances. The comparison for BC diiferentially co-expressed mRNA modules is reported in Table 10 of Supplementary file.

**Table 7:**
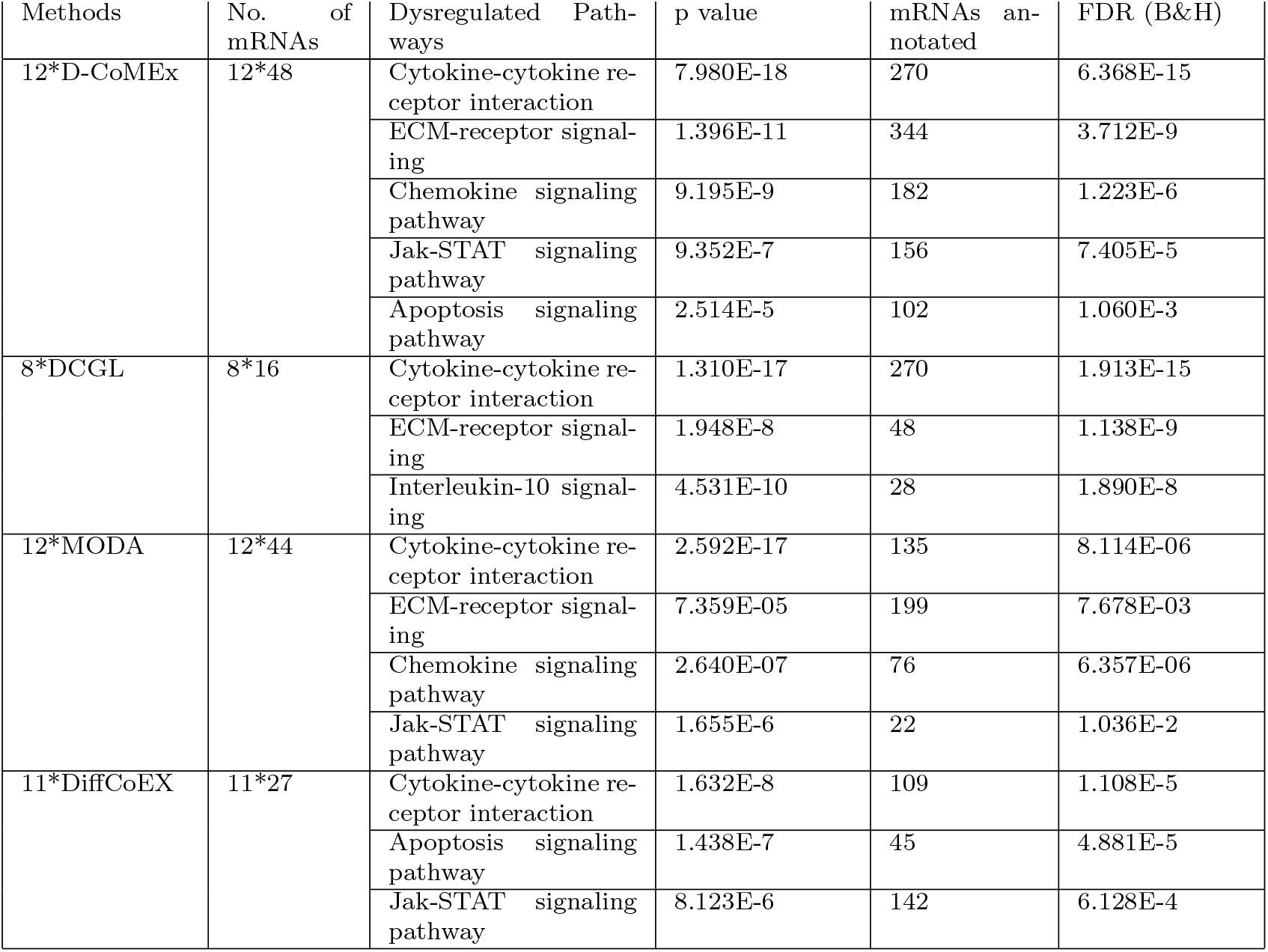
Comparison of D-CoMEx results with that of DCGL, MODA and DiffCoEx results for PD dataset.

## 4 DISCUSSION

Biomarkers which are hallmarks of a disease, which undergo perturbation in terms of topological and biological properties during progression from control (initial) stage to disease stage. In this paper, we present a novel method, which integrates a CEN technique and a differential analysis technique to extract potential miRNAs regulating dysregulated pathways from an mRNA-miRNA dataset. We use a differential co-expression analysis technique to assess the preservation of the network modules across control and disease stages.

From the mRNA and miRNA co-expressed modules, we extract interesting mRNAs and miRNAs having high semantic similarities. From these interesting mRNAs and miRNAs, we find differentially co-expressed mRNAs and miRNAs using a differential analysis technique. The differentially co-expressed modules are validated biologically and statistically to assess their GO enrichment and biological functionality. Most dysregulated pathways mapped from differentially co-expressed mRNAs or miRNAs extracted from mRNA and miRNA expression profiles are common. This establishes the fact that there is a close association between the differentially co-expressed miRNAs and the dysregulated pathways.

The differentially co-expressed modules of mRNAs and miRNAs are more statistically and biologically significant than other three methods DGCL, MODA, and DiffCoEx. Like Liu et al.[46], we find the dysregulated pathways mapped from the differentially co-expressed mRNAs and miRNAs. In addition, we find the miRNAs regulating the dysregulated pathways and assess their associations with PD and BC. The key feature of dysregulated pathways and regulating miRNAs is that both are condition specific and undergo perturbation during pathogenesis of a disease. The dysregulated pathways are further analyzed using reverse search to extract the miRNAs targeting the pathways during pathogenesis of the diseases. We experimented D-CoMEx with two mRNA-miRNA RNA-Seq disease datasets and we found that D-CoMEx is agnostic to any disease and never assumes any dependence for a specific disease. This thorough analysis of dysregulated pathways focuses on *pathway-miRNA* interaction, which is likely to aid scientists studying these specific pathways with respect to the diseases.

Based on our recommendations and in light of the above discussion, we hypothesize that there is close association between dysregulated pathways and miRNAs, and highlight the following issues for further investigation, from among many avenues for future work.

1. From dysregulated pathways identified using differential co-expression analysis, scientists can get insights into other dysregulated pathways.
2. From the differentially co-expressed miRNAs extracted from differential co-expression analysis, functionality of unexplored miRNAs in progression of a disease can be hypothesized.
3. Because of the close association between dysregulated pathways and their regulating miRNAs, discovering functionality and crosstalk of miRNAs regulating dysregulated pathways can be an interesting research topic.

## 5 CONCLUSION

The analysis introduced in this paper can be used by biologists to augment their research on biomarkers associated with diseases, and provide guidance towards effective drug development targeting biomarkers to prevent the pathogenesis of the disease.

The method we proposed is implemented using *R* and is called as D-CoMEx. The method integrates a CEN technique and a differential analysis technique to find the differentially co-expressed mRNAs and miRNAs, which are condition specific. The miRNAs, targeting the dysregulated pathways mapped from differentially co-expressed mRNA and miRNA modules of both PD and BC datasets either up-regulate or down-regulate during progression of the disease.

The proposed method can be extended to identify miRNA biomarkers involved in other diseases and to establish miRNA-disease associations, provided jointly profiled mRNA-miRNA expression datasets for those diseases are available.

## Author Contributions & Funding

All authors contributed in the design of the framework. TK implemented the method using R, and wrote the manuscript. DKB, and JK reviewed the manuscript.

There is no source of funding.

