## Supplemental for "D-CoMEx: A Unified Approach to Identify Phenotype-specific miRNA Biomarkers Related to Diseases": D_CoMEx_Supplementary.pdf

In the Supplementary file, we elaborate the preprocessing activity of the two datasets used in the experimental studies and the detailed results in order to establish the generalizability of our proposed method D-CoMEx.

- Section 1 describes the preprocessing steps of PD and BC datasets.
- In Section 2 and 3 we describe the results for dataset 1 and 2, respectively
- In Section 4, we give a detailed account for setting of parameters, namely expression similarity threshold ($\delta$) and minimum neighborhood threshold ($\rho$) through heuristic approach in Figure 1.

#### 1 Preprocessing of Datasets

In our work, we used two RNA-seq datasets with both mRNA and miRNA expression profiles to predict a strong association between miRNAs and dysregulated pathways related to both diseases. The details of the datasets are given in Table 1.

- **Dataset 1:** The jointly profiled mRNA-miRNA dataset (GSE77668) consists of 12 control and 12 disease samples for both 148 mRNAs and 822 miRNAs. Here, we normalized the gene expression values and removed batch effects using removeBatchEffect() of edgeR package [**?**].
- **Dataset 2:** We used GSE19783 (breast cancer subtypes dataset) with the following jointly profiled miRNA-mRNA expression profiles with 40996 probes across 115 samples and 489 miRNAs across 101 samples. For mRNA, we considered only two set of conditions under breast cancer subtypes, normal-like and Lum A breast cancer subtype. For normal-like, we have found 10 conditions and for Lum A, we have taken 41 conditions out of many conditions. Similarly, for miRNA, we considered only two breast cancer subtypes, with 10 normal-like conditions and 41 Lum A breast cancer subtype. We then filtered 4024 mRNAs out of 40996 which have p values $< 0.7$ using DESeq2. After mapping the 4024 probes to the gene names, we found 3714 mRNAs having gene symbol. We obtain mRNA dataset of 3714 mRNAs. Again out of 3714 mRNAs, we filtered the genes which have mapped entrezIDs with them and found 3441 mRNAs. Finally, we filtered the expression matrix for 3441 mRNAs only which have mapped entrezIDs.

Table 1: Datasets and their description

| Sl. no. | Disease | GEO ID(s) | Description |
|---|---|---|---|
| 1 | Parkinson's disease | GSE77668 | 148 mRNAs $\times 24$ samples (12 control samples and 12 disease samples)<br>822 miRNAs $\times 24$ samples (12 control samples and 12 disease samples) |
| 2 | Breast Cancer | GSE19783 | 40996 mRNAs $\times 115$ samples across multiple breast-cancer sub-types (we considered 10 control (normal) samples and 41 disease (tumor) samples)<br>489 miRNAs $\times 101$ samples across multiple breast-cancer sub-types (we considered 10 control (normal) samples and 41 disease (tumor) samples) |

Similarly, for miRNAs, we removed the miRNAs, which do not have any values and found 489 miRNAs having values.

#### 2 Analysis of D-CoMEx results using dataset 1

In Table 2, we show the association among differentially co-expressed mRNAs and miRNAs extracted from expression profiles of dataset 1 using the experimentally validated miRNA-mRNA dataset downloaded from miRWalk 2.0. Tables 3 and 4 show the enrichments of dysregulated pathways mapped from the differentially co-expressed mRNAs and miRNAs for dataset 1, respectively.

Table 2: Validation of interactions among differentially co-expressed miRNAs with the differentially co-expressed mRNAs for dataset 1

| Set of target genes from interesting miRNAs | mRNA module 1 | mRNA module 2 | mRNA module 3 | mRNA module 4 |
|---|---|---|---|---|
| Target gene set 1 | 24 | 11 | 14 | 10 |
| Target gene set 2 | 25 | 11 | 14 | 10 |
| Target gene set 3 | 28 | 12 | 15 | 11 |
| Target gene set 4 | 32 | 13 | 17 | 12 |

Table 3: Dysregulated pathways mapped from differentially co-expressed mRNAs for dataset 1

| Dysregulated Pathways | $p$ value | Dysregulated Pathways | $p$ value |
|---|---|---|---|
| Cytokine-cytokine receptor interaction | 7.98E-18 | Chemokine signaling pathway | 9.20E-09 |
| ECM receptor signalling | 1.40E-11 | Interleukin signaling pathway | 1.50E-06 |
| p53 pathway | 4.21E-05 | Apoptosis signaling pathway | 6.10E-06 |
| TGF-beta signaling pathway | 1.19E-03 | Inflammation mediated by chemokine and cytokine signaling pathway | 1.62E-05 |
| Gonadotropin-releasing hormone receptor pathway | 1.58E-03 | IL-17 signaling pathway | 3.071E-11 |

Table 4: Dysregulated pathways mapped from differentially co-expressed miRNAs for dataset 1

| Dysregulated Pathways | score | Dysregulated Pathways | score |
|---|---|---|---|
| Mapk Signaling Pathway | 1.388 | Wnt Signaling Pathway | 1.43 |
| Interleukin-10 signaling | 1.281 | T Cell Receptor Signaling Pathway | 0.946 |
| Colorectal Cancer | 0.769 | Cytokine Signaling In Immune System | 0.67 |
| Cytokines and Inflammatory Response | 0.643 | Tnf Receptor Signaling Pathway | 0.634 |
| Chemokine signaling pathway | 0.61 | p53 pathway | 0.587 |
| ECM receptor interaction | 0.571 | Lissencephaly Gene (Lis1) In Neuronal Migration And Development | 0.61 |
| Apoptosis signaling pathway | 0.515 | Parkinsons Disease | 0.57 |
| Cell Death Signalling Via Nrage Nrif And Nade | 0.479 | Cytokine-cytokine receptor interaction | 0.526 |

In Table 5, we describe common dysregulated pathways related to PD.

Table 5: Common Dysregulated pathways associated with PD

| Dysregulated Pathways | Description |
|---|---|
| Apoptosis signalling pathway | In [50], Tatton et. al have established that there is premitochitondrial apoptosis Signalling during progression of Parkinson's disease. |
| Parkinson's disease | The significant hallmark of PD is dysfunction of mitochrondria and accumulation of Lewy bodies in the neurons, due to mutations in DJ-1, PINK1, alpha-synuclein, LRRK2, and parkin, which lead to decrease in dopamine release or lead to prior death of dopaminergic neurons [?] |
| Inflammation mediated by chemokine and cytokine signalling pathway | Cytokines and chemokines are the key drivers of inflammation, hematopoiesis, and initiation of immune responses. In yesteryears, knowledge has emerged on the role of cytokines and chemokines in neurodegenerative disorders, such as PD and AD [?]. |
| Wnt signalling pathway | Wnt signalling pathway play significant role in maintaining of dopamine neurons. During pathogenesis of PD, mutation in LRRK2 and overexpression of armadillo/B-catenin alter the process of Wnt Signalling, which contribute to the progressive loss of dopamine neurons [49] |
| Integrin signalling pathway | Integrins are the key regulators of cell migration, differentiation of cells, inflammatory responses, and overall development and repair of the nervous system. Any imbalance in the integrin signalling pathway leads to neuronal imbalance and thereby play a major role in the pathogenesis of neurodegenerative disorders, such as AD and PD [?]. |
| Dopamine receptor mediated signaling pathway | Dopamine play an important role in normal brain processes, including motor behavior, memory, and loss of dopamine neurons in substantia nigra modulate the progression of neurodegenerative disease, such as, PD. [?] |
| Ionotropic glutamate receptor pathway | Dopamine and glumate interplay critical aspects during progression of PD. Loss of dopaminergic neurons alter the neurotransmission in basal ganglia motor neuron whereas glutamate receptors reverse the altered neurotransmission and debilitate the primary symptoms of PD [?] |

### 3 Analysis of D-CoMEx results using dataset 2

In Table 6, we show the association among differentially co-expressed mRNAs and miRNAs extracted from expression profiles of dataset 2 using the experimentally validated miRNA-mRNA dataset downloaded from miRWalk 2.0. Tables 7 and 8 show the enrichments of dysregulated pathways mapped from the differentially co-expressed mRNAs and miRNAs for dataset 2, respectively.

Table 6: Validation of interactions among differentially co-expressed miRNAs with the differentially co-expressed mRNAs for dataset 2

| Set of target genes from interesting miRNAs | mRNA module 1 | mRNA module 2 | mRNA module 3 | mRNA module 4 |
|---|---|---|---|---|
| Target gene set 1 | 299 | 60 | 57 | 63 |
| Target gene set 2 | 176 | 40 | 29 | 36 |
| Target gene set 3 | 172 | 44 | 34 | 34 |
| Target gene set 4 | 98 | 17 | 16 | 12 |

Table 7: Dysregulated pathways mapped from differentially co-expressed mRNAs for dataset 2

| Dysregulated Pathways | $p$ value | Dysregulated Pathways | $p$ value |
|---|---|---|---|
| Wnt signaling pathway | 1.44E-03 | Notch signaling pathway | 1.09E-04 |
| Apoptosis signaling pathway | 3.01E-03 | Cell cycle | 1.10E-04 |
| Angiogenesis | 0.00006806 | Pathways in cancer | 0.0002971 |
| Sonic Hedgehog (SHH) signaling pathway | 2.13E-05 | Blood coagulation/Hemostatis | 1.21E-04 |
| Histamine H1 receptor mediated signaling pathway | 2.15E-06 | T cell receptor Signaling pathway | 4.16E-03 |

In Table 9, we describe common dysregulated pathways related to BC.

Table 8: Dysregulated pathways mapped from differentially co-expressed miRNAs for dataset 2

| Dysregulated Pathways | score | Dysregulated Pathways | score |
|---|---|---|---|
| SHH signaling | 1.645 | Wnt Signaling Pathway | 1.59 |
| Hemostatis | 1.39 | T Cell Receptor Signaling Pathway | 1.202 |
| Angiogenesis | 3.018 | NGF Signalling | 2.44 |
| Cell cycle | 2.14 | MAPK | 3.091 |

Table 9: Common Dysregulated pathways associated with BC

| Dysregulated Pathways | Description |
|---|---|
| Cell Cycle pathway | Deregulation of cyclins D1 and E1 and the cyclin-dependent kinase inhibitor, such as INK4A play important role in cell cycle and impact severely on breast cancer [?] |
| Hemostatis pathway | Blood coagualtion is one of the important related disorder to breast cancer. In [?], author say that activation of MET oncogene promotes the invasion and metastatis in breast cancer, suggesting that an significant role of hemostatis in breast cancer. |
| WNT signaling pathway | Wnt signalling pathway play significant role in regulation of progenitor cells in human breast cancer cell lines and is sincerely involved in anticancer drug resistance [?]. |
| T-Cell receptor pathway | The presence of T cells in blood and tumorous breast suggest the existance of tumorigenesis of breast [53] |
| SHH signaling | The Sonic Hedgehog pathway is responsible for growth and proliferation of cells in mamary glands during embryogenesis. During breast cancer, there is disruptions in target genes, such as PTCH-1 and GL1-2, which lead to dysplasia of breast ducts, suggesting the significance of SHH signaling pathway in tumorigenesis of breasts [?]. |

Table 10: Comparison of D-CoMEx results with that of DCGL, MODA and DiffCoEx results for BC dataset

| Methods | No. of mRNAs | Dysregulated Pathways | p value | FDR (B&H) | mRNAs annotated |
|---|---|---|---|---|---|
| D-CoMEx | 557 | Hemostasis | 1.32E-04 | 3.81E-02 | 36 |
| | | WNT signaling pathway | 2.19E-06 | 3.05E-02 | 45 |
| | | Cell cycle | 3.15E-05 | 1.16E-03 | 56 |
| DCGL | 149 | Angiogenesis | 3.83E-05 | 2.89E-02 | 66 |
| | | Hemostasis | 8.19E-04 | 3.77E-02 | 56 |
| | | P53 pathway feedback loops | 14.16E-03 | 1.91E-02 | 60 |
| MODA | 194 | Angiogenesis | 1.01E-04 | 2.12E-01 | 11 |
| | | Cadherin signaling pathway | 2.18E-02 | 3.21E-02 | 18 |
| | | Cytoskeletal regulation by Rho GTPase | 3.09E-04 | 2.15E-02 | 31 |
| DiffCoEX | 277 | p38 MAPK pathway | 2.19E-04 | 1.2E-02 | 35 |
| | | Apoptosis signaling pathway | 7.61E-05 | 1.01E-02 | 22 |
| | | Wnt signaling pathway | 3.11E-03 | 4.14E-02 | 18 |

### 4 Significance of setting parameters in THD-Module Extractor

We use THD-Module Extractor to extract co-expressed modules from both control and disease conditions of mRNA and miRNA expression profiles. THD-Module Extractor accepts two parameters, viz., expression similarity threshold ($\delta$) and minimum neighborhood threshold ($\rho$). In our experiment, we chose expression similarity threshold ($\delta$) and minimum neighborhood threshold ($\rho$) heuristically as 0.95 and 250, respectively, throughout the experiment to optimize the results and avoid the generation of modules with less cardinality. This can be evidenced from Figure 1. The left y-axis of the graph denotes the statistical significance of the results in terms of p values, the right y-axis shows the minimum neighborhood threshold ($\rho$), and the x-axis represents the conditions. However, the threshold figure may change for different datasets depending on the number of instances, dimensionality, and data distribution.

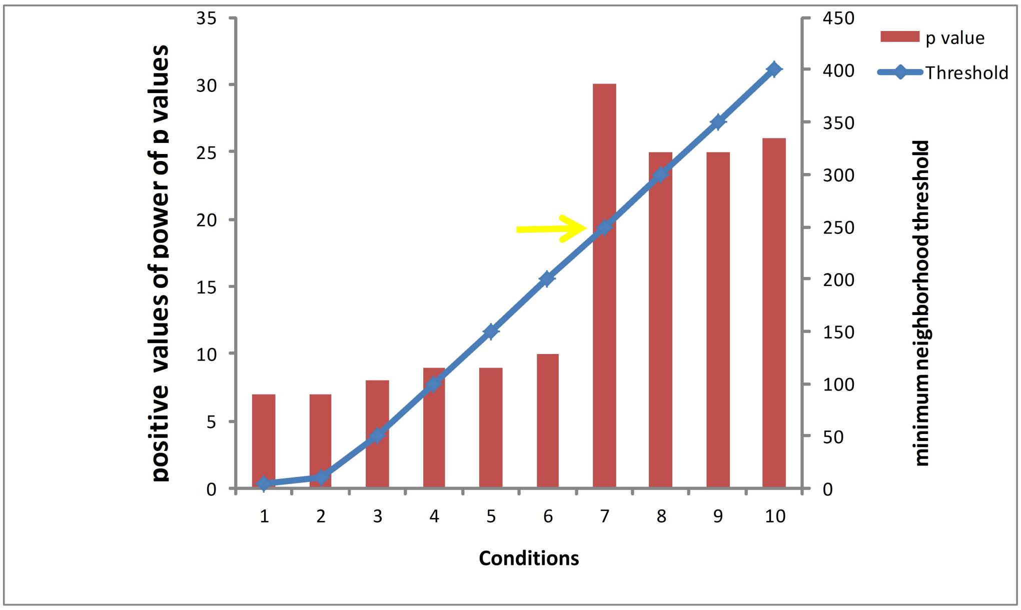


Figure 1: Minimum neighborhood threshold $\rho$ versus positive power of e of $p$ values. Here, the yellow arrow points to the minimum neighborhood threshold value (250) corresponding to the best positive power of $e$ of $p$ value (20).

[2] Tulika Kakati, Hirak Kashyap, and Dhruba K Bhattacharyya. Thd-module extractor: An application for cen module extraction and interesting gene identification for alzheimer's disease. *Scientific Reports*, 6, 2016.

[3] Ke-Qin Liu, Zhi-Ping Liu, Jin-Kao Hao, Luonan Chen, and Xing-Ming Zhao. Identifying dysregulated pathways in cancers from pathway interaction networks. *BMC bioinformatics*, 13(1):1, 2012.

[4] Sumanta Ray and Sanghamitra Bandyopadhyay. Discovering condition specific topological pattern changes in coexpression network: an application to hiv-1 progression. *IEEE/ACM transactions on computational biology and bioinformatics*, 13(6):1086–1099, 2016.

[5] Zhouxian Bai, Guangchun Han, Bin Xie, Jiajia Wang, Fuhai Song, Xing Peng, and Hongxing Lei. Alzbase: an integrative database for gene dysregulation in alzheimer's disease. *Molecular neurobiology*, 53(1):310–319, 2016.

[6] Su-Ping Deng, Lin Zhu, and De-Shuang Huang. Predicting hub genes associated with cervical cancer through gene co-expression networks. *IEEE/ACM Transactions on Computational Biology and Bioinformatics (TCBB)*, 13(1):27–35, 2016.

[7] Abolfazl Doostparast Torshizi and Linda Petzold. Sparse pathway-induced dynamic network biomarker discovery for early warning signal detection in complex diseases. *IEEE/ACM Transactions on Computational Biology and Bioinformatics*, 2017.

[8] Yuansheng Liu, Xiangxiang Zeng, Zengyou He, and Quan Zou. Inferring microrna-disease associations by random walk on a heterogeneous network with multiple data sources. *IEEE/ACM transactions on computational biology and bioinformatics*, 14(4):905–915, 2017.

[9] Duc-Hau Le. Network-based ranking methods for prediction of novel disease associated micrornas. *Computational Biology and Chemistry*, 58:139–148, 2015.

[10] Duc-Hau Le, Lieven Verbeke, Dinh-Toi Chu, Van-Huy Pham, et al. Random walks on mutual microrna-target gene interaction network improve the prediction of disease-associated micrornas. *BMC bioinformatics*, 18(1):479, 2017.

[11] Sipko van Dam, Urmo Võsa, Adriaan van der Graaf, Lude Franke, and João Pedro de Magalhães. Gene co-expression analysis for functional classification and gene–disease predictions. *Briefings in bioinformatics*, page bbw139, 2017.

[12] David Amar, Hershel Safer, and Ron Shamir. Dissection of regulatory networks that are altered in disease via differential co-expression. *PLoS Comput Biol*, 9(3):e1002955, 2013.

[13] Bruno M Tesson, Rainer Breitling, and Ritsert C Jansen. Diffcoex: a simple and sensitive method to find differentially coexpressed gene modules. *BMC Bioinformatics*, 11(1):497, 2010.

[14] Bao-Hong Liu, Hui Yu, Kang Tu, Chun Li, Yi-Xue Li, and Yuan-Yuan Li. Dcgl: an r package for identifying differentially coexpressed genes and links from gene expression microarray data. *Bioinformatics*, 26(20):2637–2638, 2010.

[15] Dong Li, James B Brown, Luisa Orsini, Zhisong Pan, Guyu Hu, and Shan He. Moda: Module differential analysis for weighted gene co-expression network. *arXiv preprint arXiv:1605.04739*, 2016.

[16] Li Zhu, Ying Ding, Cho-Yi Chen, Lin Wang, Zhiguang Huo, SungHwan Kim, Christos Sotiriou, Steffi Oesterreich, and George C Tseng. Metadcn: meta-analysis framework for differential co-expression network detection with an application in breast cancer. *Bioinformatics*, 33(8):1121–1129, 2016.

[17] Sumanta Ray and Ujjwal Maulik. Identifying differentially coexpressed module during hiv disease progression: A multiobjective approach. *Scientific reports*, 7(1):86, 2017.

[18] Darryl Nishimura. Biocarta. *Biotech Software & Internet Report: The Computer Software Journal for Scient*, 2(3):117–120, 2001.

[19] Minoru Kanehisa, Miho Furumichi, Mao Tanabe, Yoko Sato, and Kanae Morishima. Kegg: new perspectives on genomes, pathways, diseases and drugs. *Nucleic Acids Research*, 45(D1):D353–D361, 2017.

[20] Lisa Matthews, Gopal Gopinath, Marc Gillespie, Michael Caudy, David Croft, Bernard de Bono, Phani Garapati, Jill Hemish, Henning Hermjakob, Bijay Jassal, et al. Reactome knowledgebase of human biological pathways and processes. *Nucleic acids research*, 37(suppl 1):D619–D622, 2009.

[21] Jing Chen, Eric E Bardes, Bruce J Aronow, and Anil G Jegga. Toppgene suite for gene list enrichment analysis and candidate gene prioritization. *Nucleic acids research*, 37(suppl_2):W305–W311, 2009.

[28] Ioannis S Vlachos, Konstantinos Zagganas, Maria D Paraskevopoulou, Georgios Georgakilas, Dimitra Karagkouni, Thanasis Vergoulis, Theodore Dalamagas, and Artemis G Hatzigeorgiou. Diana-mirpath v3. 0: deciphering microrna function with experimental support. *Nucleic acids research*, 43(W1):W460–W466, 2015.

[29] Ron Edgar, Michael Domrachev, and Alex E Lash. Gene expression omnibus: Ncbi gene expression and hybridization array data repository. *Nucleic acids research*, 30(1):207–210, 2002.

[30] Peter Langfelder and Steve Horvath. Wgcna: an r package for weighted correlation network analysis. *BMC bioinformatics*, 9(1):559, 2008.

[31] Hasin Afzal Ahmed, Priyakshi Mahanta, Dhruba Kumar Bhattacharyya, and Jugal Kumar Kalita. Shifting-and-scaling correlation based biclustering algorithm. *IEEE/ACM transactions on computational biology and bioinformatics*, 11(6):1239–1252, 2014.

[32] Philip Resnik et al. Semantic similarity in a taxonomy: An information-based measure and its application to problems of ambiguity in natural language. *J. Artif. Intell. Res.(JAIR)*, 11:95–130, 1999.

[33] Hussain Ahmed Chowdhury, Dhruba Kumar Bhattacharyya, and Jugal K Kalita. (differential) co-expression analysis of gene expression: A survey of best practices. *IEEE/ACM transactions on computational biology and bioinformatics*, 2019.

[34] Harsh Dweep, Carsten Sticht, Priyanka Pandey, and Norbert Gretz. mirwalk–database: prediction of possible mirna binding sites by "walking" the genes of three genomes. *Journal of biomedical informatics*, 44(5):839–847, 2011.

[35] Xing Chen, Di Xie, Qi Zhao, and Zhu-Hong You. Micrornas and complex diseases: from experimental results to computational models. *Briefings in bioinformatics*, 2017.

[44] Xuan Zhuang, Zhiming Li, Huinuan Lin, Long Gu, Qing Lin, Zhongxian Lu, and Chi-Meng Tzeng. Integrated mirna and mrna expression profiling to identify mrna targets of dysregulated mirnas in non-obstructive azoospermia. *Scientific reports*, 5, 2015.

[50] William G Tatton, Ruth Chalmers-Redman, David Brown, and Nadine Tatton. Apoptosis in parkinson's disease: signals for neuronal degradation. *Annals of Neurology: Official Journal of the American Neurological Association and the Child Neurology Society*, 53(S3):S61–S72, 2003.

[51] Tien-Chun Yang, Pei-Chun Wu, I Chung, Jhih-Hang Jiang, Ming-Ji Fann, Lung-Sen Kao, et al. Cell death caused by the synergistic effects of zinc and dopamine is mediated by a stress sensor gene gadd45b–implication in the pathogenesis of parkinson's disease. *Journal of Neurochemistry*, 139(1):120–133, 2016.

[52] Alexey Koval and Vladimir L Katanaev. Dramatic dysbalancing of the wnt pathway in breast cancers. *Scientific reports*, 8(1):7329, 2018.

[53] John F Beausang, Amanda J Wheeler, Natalie H Chan, Violet R Hanft, Frederick M Dirbas, Stefanie S Jeffrey, and Stephen R Quake. T cell receptor sequencing of early-stage breast cancer tumors identifies altered clonal structure of the t cell repertoire. *Proceedings of the National Academy of Sciences*, 114(48):E10409–E10417, 2017.

[54] Bin Hao, Xin Chen, Dongwei Dai, Chao Zou, Xi Wu, and Jianchun Chen. Bioinformatic analysis of microrna expression in parkinson's disease. *Molecular medicine reports*, 11(2):1079–1084, 2015.

[55] Liang Zhao and Zhiqin Wang. Micrornas: Game changers in the regulation of $\alpha$-synuclein in parkinson's disease. *Parkinson's Disease*, 2019, 2019.

[56] YaXing Gui, Hai Liu, LiShan Zhang, Wen Lv, and XingYue Hu. Altered microrna profiles in cerebrospinal fluid exosome in parkinson disease and alzheimer disease. *Oncotarget*, 6(35):37043, 2015.

[57] Liye Zhu, Jing Gao, Kunlun Huang, Yunbo Luo, Boyang Zhang, and Wentao Xu. mir-34a screened by mirna profiling negatively regulates wnt/$\beta$-catenin signaling pathway in aflatoxin b1 induced hepatotoxicity. *Scientific reports*, 5, 2015.
